# Target-Decoy MineR for determining the biological relevance of variables in noisy data sets

**DOI:** 10.1101/2020.11.09.374181

**Authors:** Cesaré Ovando-Vázquez, Daniel Cázarez-García, Robert Winkler

## Abstract

Machine learning algorithms excavate important variables from biological *big data*. However, deciding on the biological relevance of identified variables is challenging. The addition of artificial noise, ‘decoy’ variables, to raw data, ‘target’ variables, enables calculating a false-positive rate (FPR) and a biological relevance probability (BRp) for each variable rank. These scores allow the setting of a cut-off for informative variables can be defined, depending on the required sensitivity/ specificity of a scientific question. We demonstrate the function of the *Target-Decoy MineR* (TDM) with synthetic data and with experimental metabolomics results. The Target-Decoy MineR is suitable for different types of quantitative data in tabular format. An implementation of the algorithm in R is freely available from https://bitbucket.org/cesaremov/targetdecoy_mining/.

## 1. Introduction

Extracting informative variables from a noisy background is necessary to convert massive data sets into biological models. Statistical analyses, such as the generalized linear model for non-normal data, are used to extract these relevant variables. However, setting a stringent cut-off for statistical significance leaves fewer variables for interpretation. This sensitivity-specificity dilemma (SSD) is accompanied by the difficulty to rank variables according to their biological relevance.

### 1.1 Machine learning, variable importance scoring problem

Supervised and unsupervised machine learning (ML) methods play a crucial role in dealing with the accelerated availability of big data in biology ^1^. The random forest (RF) algorithm is an ML method that builds multiple decision tree models from randomly selected variables of a massive training data set and integrates the results into a final model^2,3^. The process is computationally efficient and robust against noise. The model builder estimates an out-of-bag (OOB) error, which indicates the reliability of the model for sample classification. Computed *variable importance* values designate each variable’s contribution to the correct splitting in the decision trees ^4^. As a measure, the overall decrease of diversity by the variable expressed as the *mean decrease Gini* (MDG) is used^5,6^. Since this calculation is unbiased by the data scientist’s knowledge, RF models enable the discovery of unexpected relationships ^7^. The RF algorithm frequently serves for the variable selection and classification of genes from microarray data^8^. In mass spectrometry-based Omics, which face the same problem, i.e., large numbers of variables and relatively few experimental repetitions, data mining (DM) strategies such as RF analysis demonstrated to be superior for sample classification to clustering algorithms^9^. Thus, DM strategies are routinely employed in bioinformatics for multiclass classification and variable selection ^10^.

Despite all these RF advantages, RF lacks sparsity in the final models, representing a serious limitation when dealing with thousands of variables. Also, due to its intrinsic random nature, *variable importance* stability must be tested. Sparse partial least square discriminative analysis (sPLS-DA) is a supervised deterministic ML classification methodology performing variable selection ^10^. sPLS-DA identifies a small subset of best variables contributing to discrimination between sample classes. This subset of best discriminative variables has a variable importance score, expressed as *loadings*. Each *load* indicates the contribution to the discriminative decision function and has a direct interpretation^10^. sPLS-DA has been used to investigate single nucleotide polymorphisms (SNP) and single nucleotide variants (SNV) data^10,11^.

Both RF and sPLS-DA are powerful tools for analyzing quantitative biological data. Nevertheless, the variable selection from RF and sPLS-DA analyses remains challenging ^12^, particularly the direct interpretation of the variable importance score and setting an optimal cut-off for possibly relevant variables. One possibility for assessing the contribution of single variables (targets) is their permutation and subsequent statistical evaluation. This method performs better than previously described strategies but is computationally demanding^13^.

These ML models, RF, and sPLS-DA must be validated using an independent set to estimate their performance. Given the specificity and scarcity of metabolomic datasets, it is impossible to keep out an independent validation set and use it to get a test error^14^. Cross-validation (CV) is a straightforward and widely used methodology to estimate ML models’ prediction error^14^. k-fold cross-validation splits the data into *k* subsets. The model is trained with *k-1* subsets and validate it on the left out sub-set. Five- or ten-fold cross-validation is considered a good compromise between bias and variance ^15^. However, it depends on the number of samples available in the dataset.

In proteomics, the target-decoy (TD) strategy is commonly used for estimating the confidence of protein identifications. Spectral matching is performed against a concatenated database, which consists of correct (‘*target*’) and false (‘*decoy*’), i.e., random protein sequences. Based on the proportion of (falsely) identified decoy proteins, a false positive rate (FPR) can be calculated, assuming that target and decoy hits occur with equal likelihood^16^.

In this work, we combined the target-decoy strategy and machine learning techniques. We call this methodology the ‘*Target-Decoy MineR (TDM)*.’

## 2. Methods

### 2.1 Target-Decoy MineR algorithm

The *Target-Decoy MineR* builds a sample classification model, using a dataset of *S* samples with *T* variables (targets) (**Fig. 1a**). Decoys are generated from the targets, permuting the sample class, obtaining *D* variables. Targets and decoys are gathered to obtain a TargetDecoy data matrix (**Fig. 1b**). The *Target-Decoy MineR* provides a ranking of variables according to the variable importance, false-positive rate, and biological relevance, using the decoys as variables that are unrelated to the sample class.

**Figure 1.**
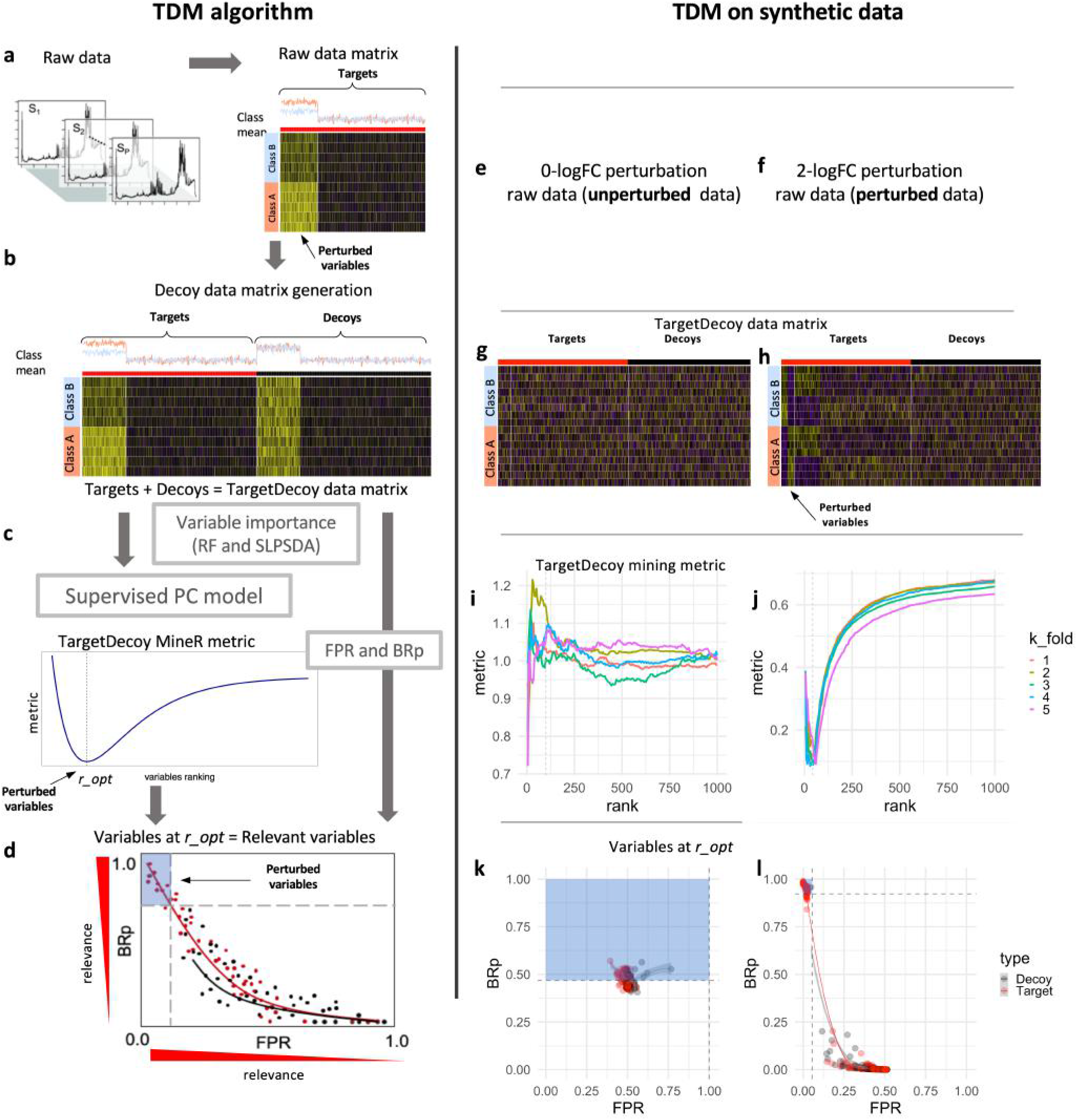
The Target-Decoy MineR (TDM) identifies biologically relevant variables in metabolomics datasets. **a-d: Algorithm**, **a** - Identification of variables in datasets representing two sample classes (A and B). **b** - Creation of a matrix with experimental (target) and artificial noise (decoy) data, **c** - Calculation of optimum metric and rank using a *k*-fold cross-validation, **d** - biological relevance probability (BRp) vs. false-positive rate (FPR) plot. **e-l: Test with synthetic data**. **e-h** synthetic data sets with 500 *m/z* variables; 50 variables are perturbed with 0- and 1-logFC. The perturbed variables correlate with the sample class, **g,h** Target-decoy matrix, **i-j** Calculation of the optimum metric and rank, using a 5-fold cross-validadion, **k-l** BRp vs. FPR plots. For the 0-logFC dataset, no relevant variables were detected (**k**); in the 1-logFC dataset, variables contributing to the classification are ranked according to their relevance (**l**).

The variable importance rank for values in the TargetDecoy data matrix is obtained by combining random forest (RF) and sPLS-DA scores. Given the variable importance ranking, a supervised principal components (sPC) model (sample classification model) with the top-ranked *m* variables is trained (**Fig. 1c**)^14^. The sPC model is generated using *C* principal components (*C ≤ S*), as input variables. The optimum model with *m* variables is obtained by varying the top-ranked variables. The minimum of the Target-Decoy MineR metric is located at the optimum variable ranking *r_opt*. At the *r_opt*, Target-Decoy MineR identifies the perturbated variables of an experiment (**Fig. 1c**), i.e., the biologically relevant variables which contribute to the differentiation of samples.

In parallel to sPC model training (**Fig. 1c**), the decoy variables enable the calculation of false-positive rates (FPR). The FPR indicates the proportion of synthetic noise variables at a certain variable importance ranking. Besides the FPR, a biological relevance probability (BRp) is calculated, using the top target and top decoy variable importance scores. The BRp is the likelihood of a variable being truly related to a sample class.

During the variable importance calculation, sPC model training, FPR, and BRp calculation, *k*-fold crossvalidation (CV) procedure is used; those validation stages prevent overfitting. In each fold, the Target-Decoy MineR trains the sPC model with all samples in the *k-1* folds and validates it with the samples in the *k* fold validation dataset. The variables in the validation dataset are pre-conditioned with the BRp ^14^.

The Target-Decoy MineR dissects the dataset and accepts or denies changing variables. Further, the variable scores (FPR and BRp) and the workflow graphics enable a straight-forward visual evaluation of the model built from the input data. Thus, the TDM allows us to judge an experiment (**Fig. 1d**) quickly. Importantly, the Target-Decoy MineR model generation takes into account variable-variable relations.

### 2.2 Synthetic data

We used the Cardinal package ^17^ to generate artificial Mass Spectrometry data using the function `simulateSpectrum()` with default parameters and `n = 5, from = 50, to = 1500, baseline = 1, by = 100`. The 5 spectra simulated were added up to obtain just only 1 spectrum. Then we used the R package `MALDIquant` ^18^ to process these spectra. First, we transformed spectra to mass spectrum profiles object using the function `createMassSpectrum` from the R package `MALDIquant`. Then, we used `thebaseline`, `alignSpectra`, `detectPeaks`, `binPeaks` with a `tolerance = 0.1, filterPeaks` with `minFrequency = 0.5`, and `intensityMatrix` functions from `MALDIquant` to get a feature matrix (simulated metabolic data). We used this methodology to create a dataset with *n* = 10 replicates and *m* = 500 variables in wild type (WT) and treatment (TR) conditions. We perturbed only 50 variables, over-expressing 25 in WT and 25 in TR. The degree of perturbation was `logFC = 0, 0.5, 1, 2.`

### 2.3 Origin and cultivation of Drosophila samples

We re-analyzed data reported by ^19^. As briefly described below and in the following sections, we grew isofemale lines of *Drosophila melanogaster* and *Drosophila mojavensis* in standard banana and yeast food at 25 °C and 12 hours light/dark. We put hundreds of flies in chambers with Petri dishes with 0.5% agar and active yeast and 5 mL of prickly pear for *Drosophila mojavensis* (to induce oviposition). We fed third instar larvae with a high protein-to-sugar or equal protein-to-sugar ratio diet^20^. We sorted adult flies by sex at 0 days of eclosion.

### 2.4 Sample preparation

We lyophilized and weighted samples. We homogenized dry-samples with a polypropylene pestle and liquid nitrogen. We added 66 μL of 85% methanol cold at −20 °C per mg of dry mass. We put samples through 5 cycles of 1-minute ultrasound at 40 kHz and 1 minute (on ice). We centrifuged samples at 15,870 g for 10 minutes at 4 °C. We collected the supernatant for *Drosophila melanogaster,* and for *Drosophila mojavensis,* we filtered samples through a 0.2 μm nylon syringe filter.

### 2.5 Non-targeted metabolic profiling

We analyzed extracts in random order for each species in distinct blocks. We used an Accela UPLC coupled to an LCQ Fleet Ion trap, using a Hypersil Gold C18 column (50 x 2.1 mm, 1.9 μm particle size) at 30 °C with a flow rate of 400 μL/min. The mobile phase was water with 0.1% (*v/v*) formic acid and methanol with 0.1% (*v/v*) formic acid. We acquired spectra in positive centroid mode in a range from 100 to 1,400 *m/z*.

### 2.6 Experimental data processing

We analyzed experimental data with `MZmine` 2.21^21^ as reported in ^19^. For feature detection, we used Grid-Mass ^22^ with a minimum relative intensity of 100. We normalized data with the maximum intensity peak. We used raw experimental data for TDM. We also removed features detected in blanks for the denoised data set before TDM.

### 2.7 Target-decoy table generation

Decoys were generated randomizing the target data (along rows and columns) using the function `sample()` from the R package `base` ^23^ until finding a population of targets and decoys with no evidence favoring the hypothesis

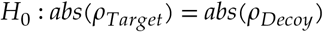

(Wilcoxon signed rank test^23^) with a significance cut-off

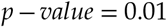

### 2.8 Univariate analysis

We used a generalized linear model (GLM) to test the hypothesis

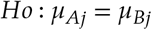

per *m/z* variable (Target or Decoy) for biological conditions *A* and *B*. We used `glm.nb()` from the R package `MASS` ^24^ with default parameters. False discovery rate (FDR) was performed with the Benjamini-Holchbech correction ^25^ function `p.adjust()` from the R package `stats`^23^.

### 2.9 Variable importance scores, multivariate analysis

We combined two-variable importance scores from two machine learning methods to get the TDM variable’s importance. These two machine learning methods are sparse partial least square discriminant analysis (SPLSDA) classifier and the random forest (RF). We developed a sparse partial least square discriminant analysis (SPLSDA) classifier model using conditions as classes and *m/z* (target and decoy) variables as features. We generated the SPLSDA with the function `splsda` from the R package `MixOmics`^10^, using default parameters for both synthetic and *Drosophila* species datasets. From the SPLSDA model, we obtained the loadings as variable importance scores. We selected SPLSDA due to its sparsity, minimizing the score of non-relevant variables. However, SPLSDA could develop low-complexity models. We developed the RF classifier models with the `functionranger()` ^26^, using the default parameters for both synthetic and *Drosophila* species datasets. We used the mean decrease Gini (MDG) values as the variable importance score. We selected RF due to its balance between model bias and complexity. The TDM variable importance of targets and decoys is the average of the mean decrease Gini (MDG) and the sparse partial least square discriminant analysis (SPLSDA) loading scores ^14^. Target and decoy variables were concatenated for calculating the target-decoy variable importance.

### 2.10 False-positive rate (FPR)

Variables were sorted in decreasing TD scores order and ranked using the `rank()` function from the R base package ^23^. We calculated FPR as the ratio of the number of decoys |*D*| to the number of decoys and targets |*D* ⋃ *T*| at rank *r*:

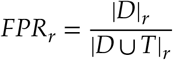

### 2.11 Target and decoy class definition

We used the top target and decoy variables (variables between the 90 and 100 percentiles) as the representative class definitions.

### 2.12 Biological relevance probability (BRp)

A support vector machine (SVM) classification model^14^ was trained using the top-ranked target and decoy variables (variables between the 90 and 100 percentiles). We used the data from the class definition mentioned previously to train the model. The SVM model delivers probabilities for all target and decoy variables, representing the biological relevance probability (BRp). A linear classification model was generated using the function `svm` from the R package`e1071`^27^ with default parameters with linear kernel, and classification type *‘C-classification.’*

### 2.13 Supervised principal components (sPC)

Top-ranked variables *m* were used to build a Principal Component Analysis (PCA) model using the R `functionprcomp`^23^. We used the *C*, `C = min(2, nrow(testData))`, principal component (PC) variables to reduce train and test data to *C* variables. We fitted a linear model on the *C* PC space using the function *lm* from the R package *stats* ^23^. The trained linear model was used to predict the class output from reduced PC validation data.

### 2.14 Pre-conditioning of data

Each variable in the validation dataset is multiplied by its corresponding BRp score, preconditioning TD variables in the validation dataset.

### 2.15 Root mean square error (RMSE)

We used the root mean square error (RMSE) to measure the sPC model prediction accuracy. We obtained validation predictions using the trained linear model and compared it to the actual validation sample class with a custom *R* function ^23^:

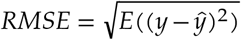

where *y* is the sample class model prediction, and 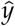 is the real sample class.

#### Metric

We defined the metric to optimize as the ratio of RMSE of predictions from the model built upon top *m* ranked variables to predictions from the model using the rest of variables (*n*) with a custom R function ^23^. The total number of variables are *total = m + n*. The TDM metric is defined as:

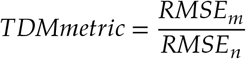

### 2.16 Cross-validation

We performed *k*-fold cross-validation using one out of *k* samples to get target-decoy scores, FPR, and BRp values. We found the optimum FPR and BRp scores in each fold, minimizing the above-mentioned metric using the *k* fold data.

### 2.17 Receiver operating curve (ROC) analysis

We used ROC analysis and Area Under the Curve (AUC) ^28^ to evaluate the global performance: sensitivity and specificity. We use the function *roc*() from the R package *pROC*^29^ to calculate AUC values.

### 2.18 Principal component analysis

The principal component analysis (PCA) TDM summary report was generated using the function `fviz_pca_var()` from the R package `factoextra`^30^.

### 2.19 Data availability statement

The data, R source, and scripts used in this study were deposited at the Bitbucket repository https://cesaremov@bitbucket.org/cesaremov/Target-Decoy_MineR.git.

## 3. Results

We tested the Target-Decoy MineR (TDM) with synthetic and experimental metabolomics datasets.

### 3.1 Processing synthetic datasets with defined perturbations

The synthetic datasets were composed of two groups, ‘wild type’ (WT) and ‘treatment’ (TR) samples, with 500 variables. Ten sample replicates were simulated, and 50 variables in each group were perturbated (25 variables in WT and 25 values TR), with *0-, 0.5-, 1-, and 2-log fold change perturbation* (**Fig. 1e-l**).

The synthetic dataset demonstrates the inverse relation between the *logFC*-fold perturbation and the TDM metric. The metric decreases as *logFC* increases (**Fig. 1i-j**). The metric approaches to zero for strong perturbations *logFC* ≥ 1. For each *logFC* perturbation, TDM produced different optimal FPR and BRp values. Optimum TDM scores are visualized in the FPR vs. BRp plots (**Fig. 1k-l**). The plots show both target and decoy points and their trend line. TDM on null perturbation data (*logFC = 0*) finds the optimal values *FPROpt = 0.49* and *BRpOpt = 0.45*, indicating the absence of discriminating variables. These values agree with the expectation that about 50% target and 50% decoy variables should be found in a dataset without two defined sample classes. Even top-ranked target variables are most likely unrelated to the sample class.

In stark contrast, if a strong perturbation was applied, such as *logFC = 2*, TDM obtained optimum values of *FPROpt = 0.00* and *BRp = 0.99*. Top-ranked variables are strongly related to the sample class. In a biological experiment, these variables are likely to be of biological relevance because they contribute to the sample differentiation.

For the intermediate case, a mild perturbation with *logFC = 0.5*, TDM obtained values of *FPROpt = 0.20* and *BRpOpt = 0.85*. Therefore, TDM detected non-random variables which are related to the sample class.

The synthetic datasets analysis demonstrated a predictable behavior of the Target-Decoy MineR results, depending on the variables’ degree of perturbations.

### 3.2 Processing experimental metabolomics data

We used experimental metabolomics data of *Drosophila melanogaster* and *Drosophila mojavensis* fed on a high protein-to-sugar ratio (HPS) or equal protein-to-sugar ratio (EPS) diet (**Fig. 2**)^19^ to test the suitability of the Target-Decoy MineR for analyzing real-life data. For investigating the influence of technical noise and the number of features, the experimental data were either analyzed ‘as is’ or after additional pre-processing of the mass spectrometry data. Raw data contained 953 features, and denoised datasets 835 features (**Fig. 2e-h**)^19^.

**Figure 2.**
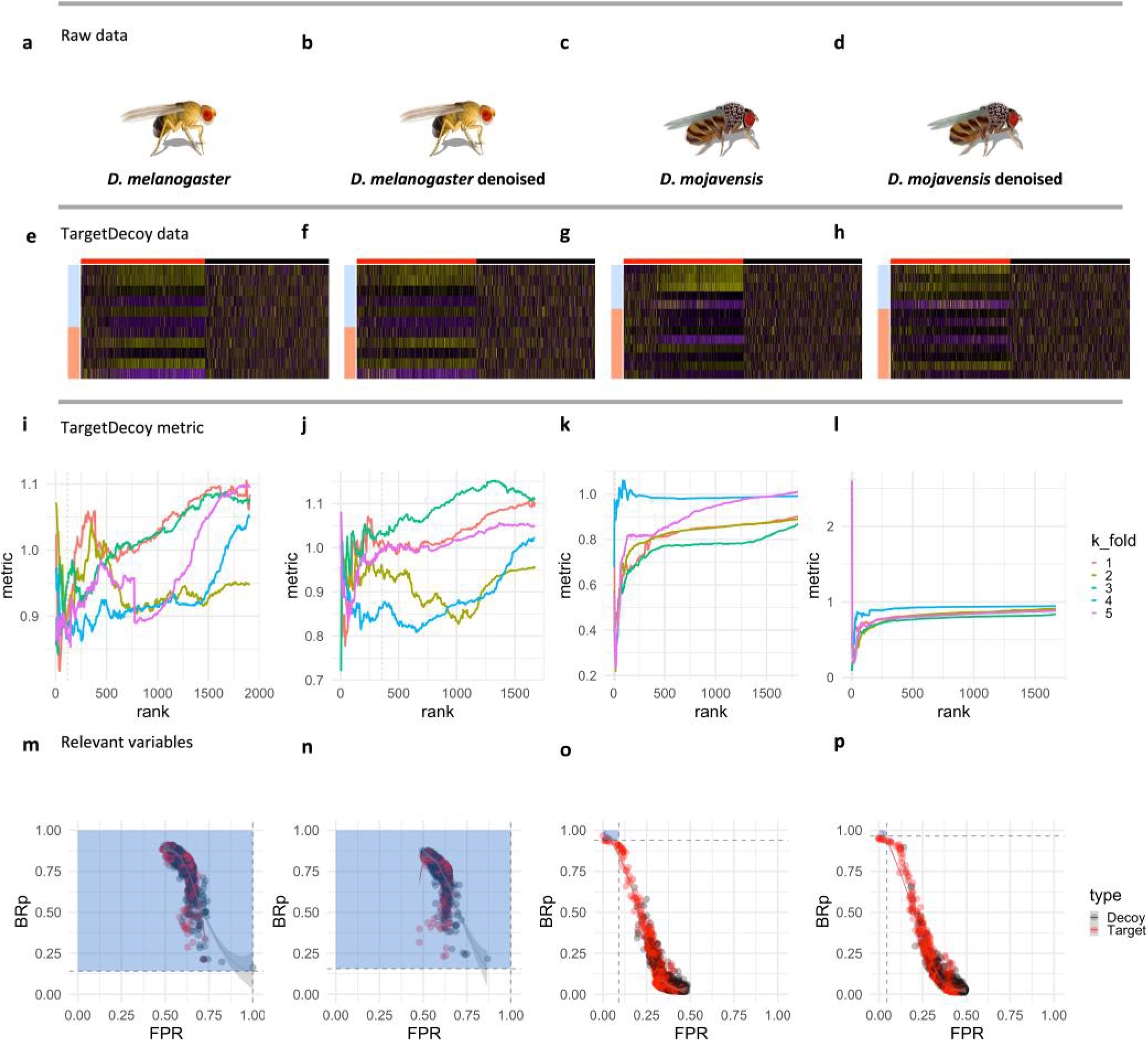
Identification of biologically relevant variables in metabolomics. *Drosophila melanogaster* and *Drosophila mojavensis* flies were fed different protein-sugar mixtures. **a-h** Target-decoy datasets obtained from raw and pre-processed (denoised) datasets. Raw datasets contain 953 targets + 953 Decoys. Denoised datasets contain 835 targets + 835 decoys, **i-l** Calculation of the optimum metric and rank, **m-p** The Target-Decoy MineR plots indicate no significant effect of the different diet on the metabolism of *Drosophila melanogaster*, but a strong effect on the *Drosophila mojavensis* metabolism. Variables which are important for the classification are not clustered together with decoy variables. The algorithm was not affected by technical noise in raw data.

The TDM analyses revealed false-positive rates (FPR) of 0.33 to 0.54 and biological relevance probabilities (BRp) of 0.50 to 0.54 for the 10 top-ranked variables of *Drosophila melanogaster*. Following their generalist and high carbohydrate diet adaptation, these values indicate the absence of a difference between the two sample classes HPS and EPS. Indeed, the two groups of flies did not exhibit a significant difference in their phenotype. In contrast, for *Drosophila mojavensis*, FPR of 0.00 and BRp values were of 0.99 to 1.00 for the top-10 variables, which indicates a stark metabolic difference between the two diets (**Fig. 2m-p**). This cactophilic fly species is highly specialized for its ecological niche and suffered developmental disorders during the experiment.

Thus, the biological effect of the treatments on the different fly species is reflected in the numerical scores of the Target-Decoy MineR.

### 3.3 Statistical performance of the Target-Decoy MineR

To investigate the statistical performance of the Target-Decoy MineR, we calculated the optimum metric and performed receiver operating curve (ROC), the area under the curve (AUC), sensitivity, and specificity calculations (**Fig. 3**). The AUC has been designated as the best single number indicator of the classification accuracy of Machine Learning algorithms^28^.

**Figure 3.**
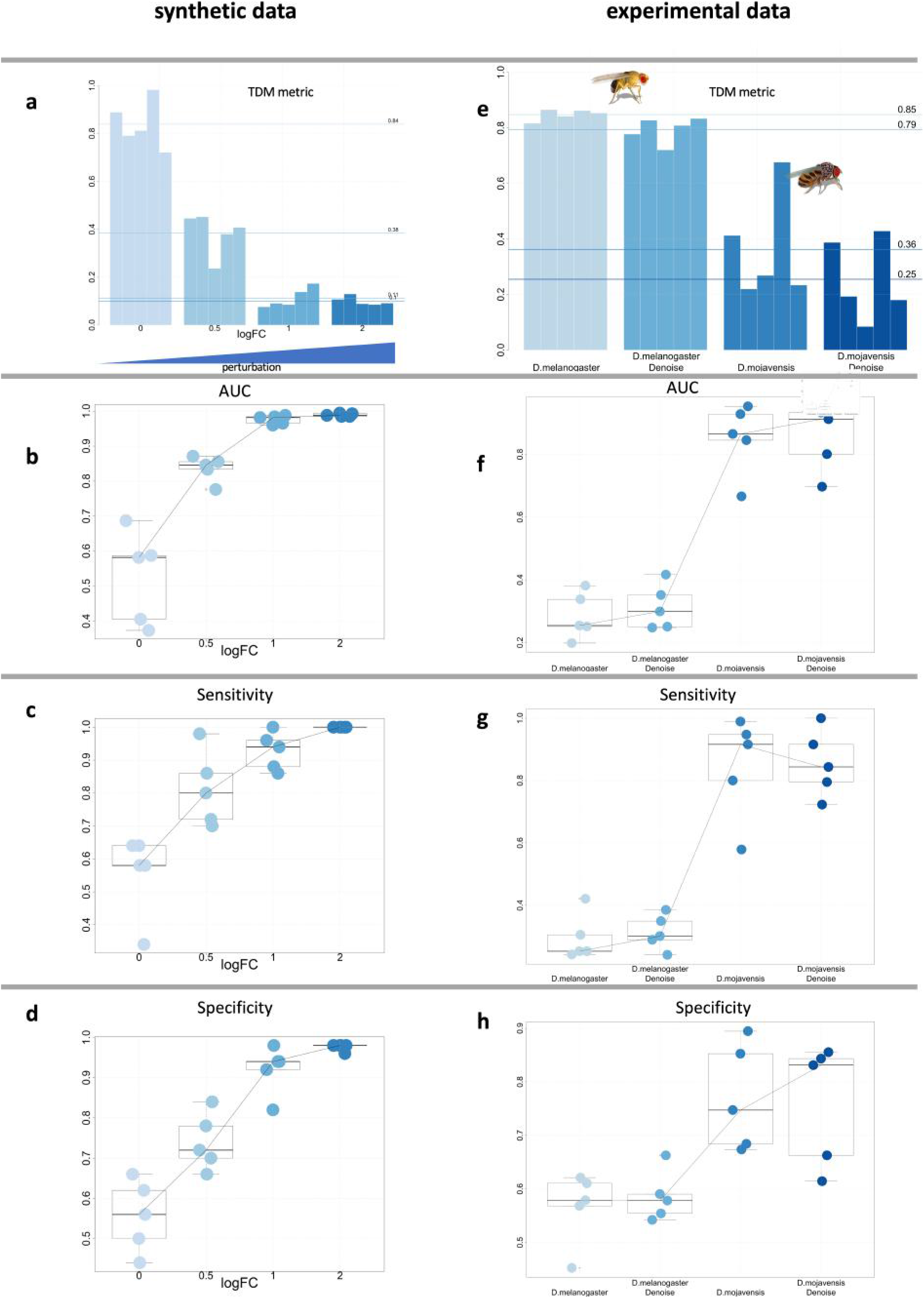
Target-Decoy MineR (TDM) evaluation of synthetic and experimental datasets. In the synthetic datasets, the TDM metric reduces as perturbation (logFC) increases. The *Drosophila melanogaster* datasets have a larger TDM metric than *Drosophila mojavensis* datasets, indicating a minor biological effect of diet on their metabolism. The area under the curve (AUC), sensitivity, and specificity increase with the system’s perturbance. The TDM worked equally well for raw and denoised metabolomics data.

For synthetic datasets with non-null perturbations (*logFC > 0*), AUC values higher than 0.6 indicate a working classifier. The AUC of the Target-Decoy MineR correlated positively with the degree of perturbation, *logFC*. A *logFC* of 0, 0.5, 1, and 2 resulted in an AUC of 0.56, 0.75, 0.94 and 0.99, respectively (**Fig. 3b**). As expected, the building of a predictive model for sample classification failed for null-perturbation data (*logFC = 0*). For data with a mild perturbation (*logFC = 0.5*) a fair model was obtained, and for *logFC > 1*, the TDM produced highly reliable models. Analogously, the sensitivity and specificity increased with the degree of the data perturbation (**Fig. 3c-d**).

Applying the Target-Decoy MineR to the *Drosophila* metabolomics data, an AUC of 0.28 was determined for *Drosophila melanogaster* and an AUC of 0.85 for *Drosophila mojavensis* (**Fig. 3f**). Sensitivity and specificity measures behave accordingly (**Fig. 3g-h**). Again, the results indicate a high tolerance of *Drosophila melanogaster* flies against a change of their diet, equally tolerated both diets. In contrast, an accurate predictive model was created for *Drosophila mojavensis*, indicating that *Drosophila mojavensis* had metabolic changes.

### 3.4 Robustness of the Target-Decoy MineR against noise

We evaluated the TDM algorithm’s sensitivity against noise comparing TDM results from using raw and denoised *Drosophila* data (**Fig. 2** **and** **3**). To denoise the data, we removed *m/z* variables below a signal-to-noise threshold^18^. For *Drosophila melanogaster*, TDM obtained top-10 BRp scores ranging from 0.33 to 0.54 for raw data and top-10 BRp scores between 0.44 to 0.57 for denoised data. Both results are similar and indicate a negligible effect of diet on the metabolic profile of *Drosophila melanogaster*. Contrary, for *Drosophila mojavensis*, TDM determined BRp scores of 1.00 for the top-10 variables of both raw and denoised data, indicating a strong biological effect. These top-10 variables are 703.67, 702.70, 704.67, 683.74, 677.68, 676.72, 635.61, 633.76, 505.95, and 502.71 *m/z*, and could be related to phospholipids with long and saturated fatty acids, which are associated with metabolic diseases and physiological desert life adaptation^19^. Using raw or pre-processed data had no significant impact on the outcome of the Target-Decoy MineR.

## 4. Discussion

The Target-Decoy MineR (TDM) finds the optimal false-positive rate (FPR) and biological relevance probability (BRp) values by minimizing the metric of variable ranks (see **Fig. 1**). This metric compares a model built using ‘informative variables’ *m* (top-ranked) to a model built upon ‘uninformative’ variables *n*. This metric can be thought of as an ‘information loss’ parameter. Thus, the selected model minimizes the ‘sample class information loss’ of variables ((**Fig. 1** **and** **3a,b**)).

For synthetic data, usable predictive models (AUC = 0.75) were created for even mildly perturbated (*logFC = 0.5*) variables. Accurate classification models with an AUC higher than 0.9 were possible with perturbations *logFC > 1*. The AUC, sensitivity, and specificity increase with the system’s perturbance for synthetic and experimental data (**Fig. 3**).

A Target-Decoy MineR model for raw and preprocessed metabolomics data of *Drosophila mojavensis* yielded 80 and 70 optimal variables, respectively. The correlation between raw and denoised BRp scores was 0.80, with a *p*-value of 0.0001. This result underlines the utility of the Target-Decoy MineR for unprocessed experimental data.

## 5. Conclusion

The addition of artificial data noise variables to experimental data further enhances the utility of machine learning strategies. The computed false-positive rates (FPR) and biological relevance probabilities (BRp) facilitate the separation between informative and uninformative variables. The Target-Decoy MineR (TDM) identifies biologically relevant variables from noisy data, even for poorly designed experiments with low statistical power, e.g., with few replicates and mild perturbation. The Target-Decoy MineR can be employed for experimental data classification and hypothesis testing.

Whereas the specificity-sensitivity dilemma is not entirely solved, now an informed decision for a reasonable cut-off for ranked variables is possible, using the FPR and BRp values. For example, in exploratory Omics studies, relatively high FPR and low BRp cut-offs could be acceptable, whereas, in detection biomarker search, strict criteria must be applied.

Strikingly, the Target-Decoy MineR is insensitive to data noise and applicable to pre-processed and raw experimental data.

The Target-Decoy MineR is easy to adapt to diverse types of massive data. An implementation of the Target-Decoy Miner as an R package is available at https://bitbucket.org/cesaremov/targetdecoy_mining/, released under the GNU General Public License (GPL). The program is immediately applicable for tabular data.

## 6. Acknowledgments

The work was funded by the CONACyT-DFG grant 2016/277850 to RW. DCG acknowledges CONACyT for his scholarship. COV acknowledges the Cátedras CONACyT program.

Authors acknowledge IPICYT Supercomputing National Center (CNS) Thubat Kaal 2.0 computing resources provided to test Target-Decoy MineR under the project TKII-R2018-COV1.

## 7. Author contributions

DCG performed biological experiments. COV, DGC, and RW designed the study, analyzed the data, and wrote the manuscript. COV programmed and tested the *Target-Decoy_MineR R package*.

